# Molecular physiology of Antarctic diatom natural assemblages reveals multiple strategies contributing to their ecological success

**DOI:** 10.1101/2023.04.19.537459

**Authors:** Carly M. Moreno, Maggie Bernish, Meredith G. Meyer, Zuchuan Li, Nicole Waite, Natalie R. Cohen, Oscar Schofield, Adrian Marchetti

**Affiliations:** Department of Earth, Marine and Environmental Sciences, University of North Carolina at Chapel Hill, Chapel Hill, NC 27599, USA; Skidaway Institute of Oceanography, University of Georgia, Savannah, GA 31411, USA; Division of Natural and Applied Science, Duke Kunshan University, Suzhou, Jiangsu 215316, China; Department of Marine and Coastal Sciences, Rutgers University, New Brunswick, NJ 08901, USA

**Author notes:** Present Address: Marine Microbiomics Laboratory, Biology Program, New York University Abu Dhabi, Abu Dhabi 129188, United Arab Emirates (Carly M. Moreno). Corresponding author. Adrian Marchetti. **Author Contributions:** CMM and AM designed the study. CMM and MB performed research. MGM, ZL, NW contributed to data analysis. NC supported sequencing analysis. OS supported research. CMM and AM wrote the manuscript. **Competing Interest Statement:** The authors disclose no competing interests.

**Keywords:** phytoplankton, diatoms, polar, Antarctica, metatranscriptomes

## Abstract

The continental shelf of the Western Antarctic Peninsula (WAP) is a highly variable system characterized by strong cross-shelf gradients, rapid regional change and large blooms of phytoplankton, notably diatoms. Rapid environmental changes coincide with shifts in plankton community composition and productivity, food web dynamics and biogeochemistry. Despite progress in identifying important environmental factors influencing plankton community composition in the WAP, the molecular basis for their survival in this oceanic region, as well as variations in species abundance, metabolism and distribution remain largely unresolved. Across a gradient of physicochemical parameters, we analyzed the metabolic profiles of phytoplankton as assessed through metatranscriptomic sequencing. Distinct phytoplankton communities and metabolisms closely mirrored the strong gradients in oceanographic parameters that existed from coastal to offshore regions. Diatoms were abundant in coastal, southern regions, where colder and fresher waters were conducive to a bloom of the centric diatom, *Actinocyclus*. Members of this genus invested heavily in growth and energy production; carbohydrate, amino acid and nucleotide biosynthesis pathways; and coping with oxidative stress, resulting in uniquely expressed metabolic profiles compared to other diatoms. We observed strong molecular evidence for iron limitation in shelf and slope regions of the WAP, where diatoms in these regions employed iron-starved induced proteins, a geranylgeranyl reductase, aquaporins, and urease, among other strategies, while limiting the use of iron-containing proteins. The metatranscriptomic survey performed here revealed functional differences in diatom communities and provides further insight into the environmental factors influencing the growth of diatoms and their predicted response to changes in ocean conditions.

## Introduction

The Western Antarctic Peninsula (WAP) is surrounded by a highly productive marine environment where phytoplankton such as large diatoms form the base of a rich polar marine food web (1–3). Diatoms are one of the main contributors to large phytoplankton blooms and carbon fluxes that occur along the coast and sea ice edge of the WAP (4, 5). Certain diatoms are efficient vectors of carbon export to the deep ocean due to their large size and heavily silicified frustules, along with efficient grazing by their euphausiid predators (6–8). However, the WAP has been undergoing rapid climate change with substantial wintertime atmospheric warming since the 1950’s (9), drastic decreases in the duration of seasonal sea-ice coverage (10), and accelerated retreat and melting of glaciers (11).

Over 28 years of sampling as part of the Palmer Long Term Ecological Research (Pal-LTER) Project has revealed factors that are critical in explaining phytoplankton dynamics and triggering of blooms in the WAP (4, 12, 13). These include strengthening of winds over the Southern Ocean, an increasing positive trend in Southern Annular Mode (SAM), and ocean warming from incursions of warm Circumpolar Deep Water (CDW) onto the shelf (9, 14), all of which control upper ocean stability, thus influencing nutrient and light availability to phytoplankton.

Although the WAP is a highly variable system and warming has plateaued in recent years (14), regional warming persists, resulting in a latitudinal climate gradient in which a maritime subpolar climate in the north transitions to a dry and cool polar climate in the south, differentially affecting phytoplankton community composition and productivity (15). In the northern region of the WAP, declines in sea ice and increased winds have resulted in deeper mixed layers and decreases in mean light levels in the upper water column (12), causing a significant reduction in summertime surface chlorophyll concentrations and more frequent occurrences of cryptophytes dominating the phytoplankton community (3). While cryptophytes have been detected throughout the entire WAP region (16), their tolerance for lower salinities (17) means they are more often associated with meltwater, and do not typically co-occur with diatoms (18). The shift from diatom biomass to small cryptophytes could favor a transition in zooplankton grazers to salps because they are more efficient than krill at grazing small cells (3, 17), potentially resulting in major shifts in the distribution of krill and the higher trophic levels they sustain. In the southern region of the WAP, there is an increase in surface chlorophyll concentrations due to the reduction in permanently ice-covered regions, allowing more light to reach phytoplankton residing under the ice and fostering blooms of large centric diatoms (2), potentially resulting in long-term shifts in the phytoplankton community (19).

Large regions of the pelagic Southern Ocean are iron (Fe) limited. Natural and artificial Fe fertilization experiments in the region have shown that resident diatoms are Fe-limited and that both pennate diatoms, such as *Fragilariopsis*, and centric diatoms, such as *Chaetoceros* and *Eucampia* (20, 21), are main responders to Fe inputs. Despite plentiful micro- and macronutients in the coastal waters of the WAP, there exists a cross-shelf gradient in Fe concentrations with a possibility of transient Fe limitation offshore where Fe-laden glacial meltwater has less reach and influence (4, 22, 23), thus creating an Fe mosaic in the region. In these regions, phytoplankton are believed to have evolved specific molecular mechanisms to cope with low Fe availability (24).

In culture and in natural populations, diatoms experiencing low Fe will frequently activate a common suite of iron mitigation strategies including luxury Fe storage (25, 26), high affinity Fe concentrating and transport mechanisms (27–29), and substitution of Fe-containing proteins with Fe-free equivalents (30, 31). A switch between Fe-containing proteins and ones containing other metal cofactor appears to be a permanent adaptation in some Antarctic diatoms (24). Light-driven proteorhodopsins have also been proposed to be useful under Fe limitation in oceanic (26, 32) and polar diatoms, where they may be involved in an alternative form of energy production to photosynthesis. Antarctic polar diatoms on the shelf and slope regions of the WAP likely employ a suite of these molecular strategies, coupled with photophysiological adaptation and acclimation strategies to reduce Fe demands in this variable light environment (36, 37).

Numerous studies along the WAP have aided our understanding of the effects of changing environmental factors on phytoplankton community composition and structure through the use of high-performance liquid chromatography (HPLC) (38) and the 18S rRNA gene marker (8, 39). For example, Luria et al. (39) examined 18S rRNA genes from water samples collected at the four corners of the Pal-LTER sampling grid and detected a 2-fold higher species richness in eukaryotic plankton in the northern WAP compared to the southern WAP. In Northern Marguerite Bay, a 15-year time series demonstrated that in years characterized as having high phytoplankton biomass, large pennate diatoms (>20 μm) dominated the 18S rRNA gene community in early spring (December), followed by large centric diatoms in summer (January) (40, 41). Further linking phytoplankton composition to biogeochemistry, Lin et al. (8) conducted high-throughput sequencing of the 18S rRNA gene and determined that out of a diverse plankton community of approximately 464 identified operational taxonomic units (OTUs) along the WAP, only a few OTUs were correlated with the spatial variability in carbon export potential.

In contrast, only a few studies have implemented gene expression analyses to investigate how polar diatom communities are able to subsist and thrive in the Southern Ocean. Metatranscriptome studies in the Ross Sea have demonstrated the potential for co-limitation between Fe and cobalamin (vitamin B_12_) (42) and the interactive effects of Fe and ocean warming on the metabolism of *Fragilariopsis* and *Pseudo-nitzschia* (43). In the northern Bransfeld Strait (north of Palmer Station and the LTER sampling grid), metatranscriptome analysis demonstrated that light-limited, sea-ice communities expressed depleted carbohydrate and energy metabolism and increased light energy dissipation, suggesting either irradiance stress or inorganic C limitation (44). These studies provide a detailed understanding of diatom metabolism in natural and nutrient-amended assemblages; however, the metabolic priorities and Fe-related gene expression patterns of diatoms have not been confirmed in the WAP. Furthermore, a cross-shelf metabolic analysis of this dynamic region, characterized by strong geochemical gradients, has not been performed and will contribute to our understanding of how this critical marine ecosystem will change under future climate change scenarios.

To better understand how environmental parameters such as Fe availability in the WAP influence the community structure and metabolism of phytoplankton, particularly that of diatoms, a metatranscriptome sequencing analysis was performed along the Pal-LTER sampling grid on samples collected in 2018. Genes involved in nutrient acquisition, photosynthesis and Fe homeostasis in diatoms were examined, providing evidence for alternative metabolic strategies among ecologically important diatoms in coping with changes in environmental conditions along the cross-shelf and north-south gradients. In addition, we opportunistically sampled a large *Actinocyclus* bloom in southern coastal sites of the Pal-LTER during our sampling period and provide a unique examination of the bloom through gene expression analysis.

## Results and Discussion

Despite some progress in identifying the important environmental factors influencing plankton community composition and function in the WAP region (8, 41, 43, 44) the molecular basis for variations in species abundance and distribution throughout the region remain largely unresolved. Here we identify oceanographic variables that influence eukaryotic phytoplankton composition in the surface waters of the WAP region. In addition, we examined the metabolic profiles of phytoplankton with an emphasis on expression patterns of iron-related genes in diatoms as assessed through metatranscriptomic sequencing. Few studies have implemented metatranscriptomic approaches in the Southern Ocean. By leveraging recently sequenced transcriptomes from ecologically relevant polar species (e.g. 23), we sought to characterize the diversity and taxon-specific metabolic activities of WAP phytoplankton.

### Oceanographic environment across the Western Antarctic Peninsula region

During the 2018 Pal-LTER cruise, sea surface temperatures and salinity decreased gradually from north to south along the WAP, with fresher waters in coastal regions relative to those over the shelf or slope, reflecting the effects of sea ice and melt waters on water column structure (Supplemental Fig. 1A, Supplemental Table 1). The northern coastal region had the warmest and saltiest waters, indicating this region was less influenced by glacial or sea ice melt and possibly more influenced by warm Upper Circumpolar Deep Water (UCDW).

Historically, there is high spatiotemporal variability in oceanographic conditions in the region although strong correlations between parameters are still observed (15, 38). Sea surface temperature and salinity were significantly positively correlated, along with dissolved inorganic carbon (DIC) and nitrate/nitrite concentrations. Chl *a*, a measure of phytoplankton biomass, was significantly correlated with both primary productivity (PP) and bacterial productivity (BP), however the two productivity rates were not significantly correlated to each other (Fig. 1B). Although not significantly correlated in 2018, a long-term study of bacterial production in the WAP found statistically significant relationships among these three parameters (15). Further, while the BP/PP ratio is low in the WAP relative to the global ocean (50), BP can at times be quite high (51); however, the variability in these relationships makes it difficult to predict BP from either PP or Chl *a* from one year to the next. Finally, in 2018, we observed a strong negative correlation between Chl *a* (and to a lesser extent PP and BP) and salinity, DIC, phosphate and nitrate(+nitrite) concentrations, indicating a drawdown of nutrients by more abundant, actively growing phytoplankton in fresher surface waters. These results are consistent with findings that high phytoplankton biomass, particularly that of diatoms, is responsible for seasonal nutrient depletions (15, 52).

**Figure 1.**
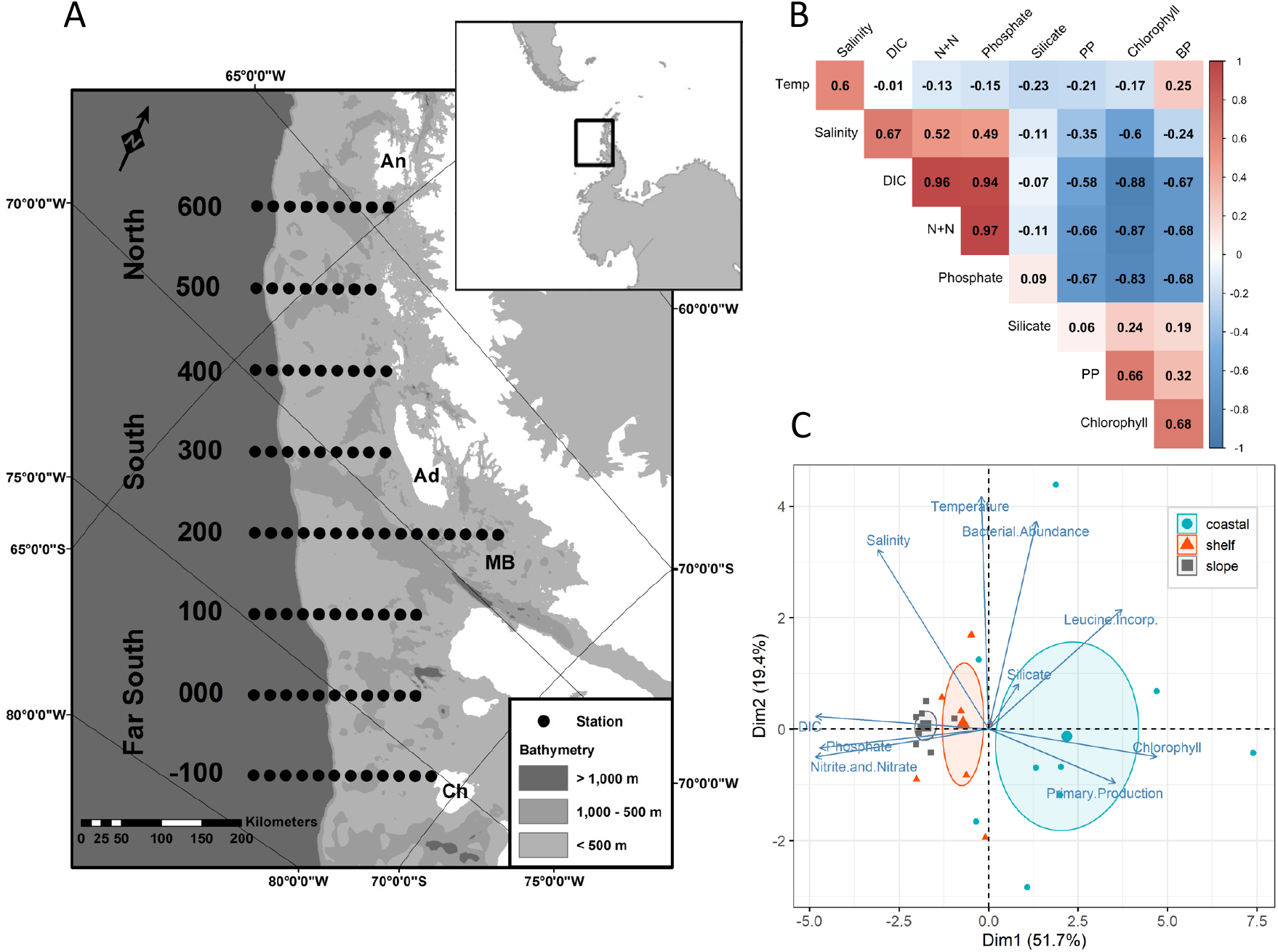
A) Location of the Palmer LTER sampling grid along the Western Antarctic Peninsula – Adapted from Conroy et al. 2020. B) Pearson’s correlation between environmental parameters. C) Principal component analysis (PCA) bi-plot of environmental measurements in surface waters, with stations colored according to coastal, shelf and slope regions.

As many of the physicochemical parameters measured were strongly correlated (either positively or negatively) with each other, we performed a principal component analysis (PCA) to examine the differences in oceanographic variables across the different sampling sites, categorized into either a) coastal, slope or shelf stations (Fig.1C) or b) North, South or Far South stations (Supplemental 1B). The first dimension (cross-shelf variability) explained 51.7% of the variance among samples and is represented by the strong negative correlation between nutrient concentrations and phytoplankton biomass (i.e., Chl *a*) that existed along a coastal to offshore gradient. Although nutrient concentrations were lower in coastal regions due to nutrient depletion by phytoplankton, fueling increased primary productivity, at no station was nitrate(+nitrite) concentration deemed to be fully depleted and thus limiting to phytoplankton growth (> 15 µmol L^−1^) (e.g., Supplemental Fig. 1A). All surface macronutrients (i.e., nitrate, silicic acid and phosphate) are generally found to be in relatively high concentrations in the Southern Ocean. This is primarily a result of there being high nutrient concentrations in the deep waters, deep winter mixing from the Upper Circumpolar Deep Water (UCDW) that resupplies the surface layer following biological depletion, and limitation of phytoplankton growth by other resources (e.g., Fe and light) (22, 53).

The second dimension (along-shelf variability) explained 19.4% of the variation among samples and was represented by oceanographic parameters that varied along a north to south gradient (Supplemental Fig.1B). Bacterial abundances were found to be higher in northern coastal regions, where temperature and salinity were also elevated (Supplemental Table 1). Previous work has shown that temperature is not a good predictor of bacterial abundance in this region and that bacterial abundances do not vary widely spatiotemporally (22); however, the northern coastal stations in 2018 had over an order of magnitude greater bacterial abundance than commonly observed in the region (14). Latitudinally varying influences on phytoplankton productivity or abundance appear to be minimal. Thus, in relation to factors influencing phytoplankton, the spatial cross-shelf differences were more significant than those present from north to south.

### Metatranscriptome sequencing statistics

To examine the diversity and inferred metabolic activity of phytoplankton communities of the WAP region, 25 samples collected from sites throughout the 2018 Pal-LTER cruise were sequenced to a depth of 13-15 million paired-end reads per sample (Supplemental Table 2). Following sequence quality control and assembly, over 15 million contigs were generated, with an N50 of 386, and combined into a large consensus assembly. Annotation of the consensus library for gene function (KEGG KO) and taxonomic association (PhyloDB) resulted in 2.3 million contigs with KEGG-annotations and 7.6 million contigs with taxonomic annotations. Of these, 38% of the KO annotations had an associated module (MO) level annotation, which links the assigned functional gene to a broader level KEGG category or pathway.

### Eukaryotic plankton community composition and diversity

Metatranscriptome assembly and taxonomic annotation of sequence read counts to assembled contigs revealed that over the entire WAP region, dinoflagellates (37.9%), diatoms (23.1%), haptophytes (21%), and cryptophytes (2.7%) constituted the dominant taxonomic plankton groups among the transcript pool (Fig. 2A; Supplemental Fig. 2). Samples from the northern sites (line 400 – 600; Fig. 1A) were predominantly comprised of transcripts assigned to dinoflagellates, while those from the far southern coastal stations (−100 – 100) had proportionally more diatom sequences, the majority of which were attributed to pennate diatoms (14.6%) rather than centric diatoms (4.7%). Northern stations were also characterized as having more diatoms in shelf water sites, while far south sites also contained more diatoms on the coast. Shannon-Weiner diversity of the eukaryotic plankton community was not significantly different for either onshore to offshore or North to Far South regions (Supplemental Fig. 3A).

**Figure 2.**
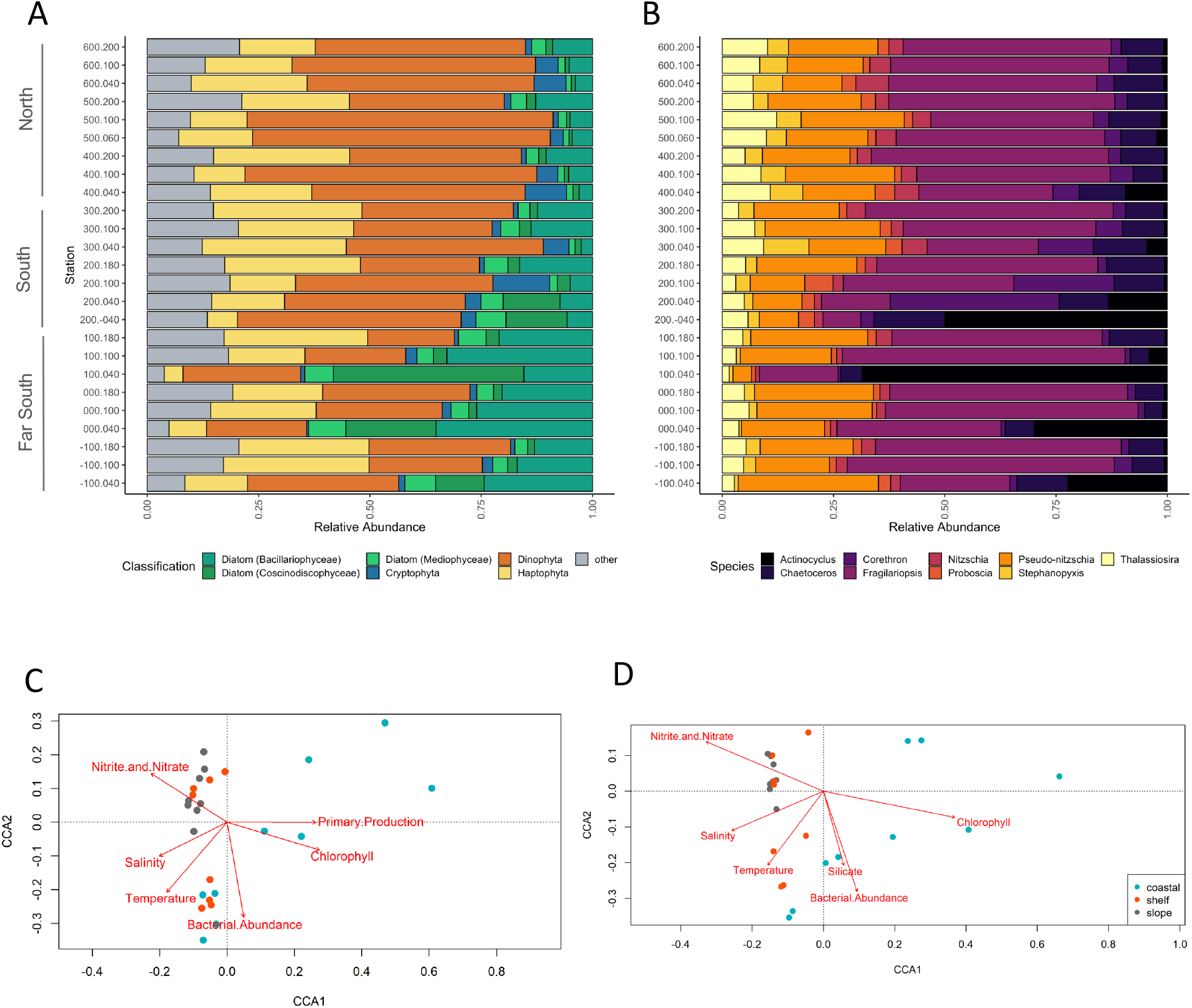
A) Taxonomic proportion of transcripts recruiting to the top most abundant phytoplankton groups. Diatoms are further delineated into pennates, radial and bi-multi polar centrics. Grey indicates all other taxonomic annotations. B) Taxonomic annotations of the top 10 most abundant diatom groups. Canonical correspondence analysis (CCA) biplots of C) the entire phytoplankton community and D) the diatom community at each station colored by distance from shore. Environmental variables are indicated by red vectors.

Of the diatoms, transcripts assigned to the genus *Fragilariopsis* were numerically dominant over the majority of sites, especially those on the shelf and slope, reaching 62.5% of transcripts at some sites. In contrast, *Actinocyclus*, a large centric diatom, constituted < 1% of communities in several northern sites, yet in several Far South sites (stations 100.040 and 000.040 specifically) comprised up to 74% of all eukaryotic transcripts, aligned to the transcriptome of an isolate of *Actinocyclus actinochilus* (UNC1410) previously sequenced from the WAP region. Additionally, station 100.040 had the highest primary production and Chl *a* measured during the cruise (Supplemental Table 1), inferring a large bloom of *Actinocyclus* in the region (Fig. 2B).

Diatom diversity significantly increased northward, indicating that non-bloom stations were more diverse, i.e., not dominated by a single or few blooming genus/species (Supplemental Fig. 3B). *Pseudo-nitzschia* constituted < 32% of the diatom relative abundance at all stations, although up to 60% of *Pseudo-nitzschia* transcripts at some stations were assigned to *P. subcurvata* at the species level, which was previously isolated from the WAP region and has a comprehensive transcriptome available (24). Other ecologically dominant centric diatoms such as those belonging to the genera *Thalassiosira*, *Chaetoceros*, *Corethron* and *Proboscia* were detected but typically comprised < 10% of the diatom community.

Higher abundances of centric diatoms have been observed using several identification methods in the WAP region. Previous 18S rRNA gene sequencing revealed large blooms of *Stellerima* and *Proboscia* in regions around Palmer Station (600 line) (8) and Marguerite Bay (200 line) (40). Taxonomically annotated mRNA transcripts from off the Wilkins Ice Shelf (−100 line) identified communities dominated by *Fragilariopsis* (44). At Rothera Station (200 line), seasonal analyses of plankton community composition measured by HPLC (40) and light and scanning electron microscopy (54) documented that the dominance of certain phytoplankton groups consistently corresponded to certain phases of seasonal progression. The phytoplankton community transitioned through three phases: from high biomass of *Phaeocystis* in early spring when wind-driven mixed layer depths (MLD) were still deep with low overall light levels, to a pennate diatom community dominated by *Fragilariopsis* as MLDs shoaled and light levels increased, and finally to a centric diatom community dominated by *Proboscia* in late summer when increased ice melt water promoted a well-stratified cold surface layer. During our collection period, the dominance of *Actinocyclus* in the coastal southern regions suggests there may be a mid-to late summer bloom pattern associated with several different diatom genera (Fig. 4). However, transcript abundance may vary in relation to physiological status of the organisms and the ratio of RNA:DNA by taxon (55, 56). For example, cryptophytes have been shown to contribute up to 50% of Chl *a* concentrations in this region (16), yet cryptophyte-assigned reads were not readily detected in our sequence libraries, generally comprising < 2% of overall transcript abundance.

Cryptophytes may not be as transcriptionally active as diatoms or dinoflagellates, which could account for their low relative transcript abundances, or there may be differences in mRNA recovery or representative database sequences (57). Inferred low cryptophyte cell abundances may also be the result of niche exclusion by dinoflagellates and diatoms as these two groups generally grow faster and can inhabit a wider range of temperatures and salinities (38, 41). Notwithstanding clear niche differentiation between these three groups, and high interannual variability in environmental parameters, taxonomic annotations derived from metatranscriptomes and the inferred eukaryotic plankton composition patterns should be interpreted with caution.

Canonical Correspondence Analysis (CCA) was used to investigate how plankton community structure as inferred from mRNA sequence libraries correlated with specific oceanographic variables. Broadly, community eukaryotic plankton composition based on metatranscriptomic read counts reflected the regions from which the communities originated (Fig. 2C). Coastal communities were associated with waters containing more Chl *a* and higher PP. Slope and shelf communities clustered into two groups, one correlated with higher temperatures and salinities in the North, and the other correlated with higher nutrient concentrations. The physicochemical variables measured explain approximately 60% of the variance in community composition (CCA1 & 2), (PERMANOVA p < 0.001). Although bacterial production was weakly significantly correlated (p = 0.05) with eukaryotic plankton community structure, it is likely not a determinant, as previously discussed.

Inferred diatom communities were further segregated according to distance from shore (Fig. 2D). The first canonical root, explaining most of the diatom variation (48.1%), revealed a notable separation between coastal and offshore stations. The strongest predictors were temperature and silicic acid concentrations (*p* = 0.01) followed by nitrate/nitrite concentrations (*p* = 0.04), with all physicochemical variables explaining 71% of variance in the diatom communities (PERMANOVA p < 0.001). Overall, temperature, PP and Chl *a* are the primary variables explaining transcript-derived diatom community composition. Other studies in this region demonstrated similar findings using either 18S rRNA gene sequencing or HPLC. In a decadal study of phytoplankton pigments along the WAP, Schofield et al. (36) found the environmental factors that favored a shallower MLD resulted in larger blooms of diatoms compared to other taxa. In a five-year study, Lin et al. (19) observed high relative abundance of centric diatoms in waters characterized by colder temperatures, reduced macronutrient concentrations, and shallow MLD, indicative of a strong influence of sea ice melt (18). Furthermore, they suggested that keystone diatoms taxa such as *Thalassiosira*, *Odontella*, *Porosira*, *Actinocyclus*, *Proboscia* and *Chaetoceros* were primarily responsible for high NCP and associated C export potential from the mixed layer.

### Variations in inferred diatom metabolism across WAP communities

To gain an understanding of diatom metabolism, we first examined broad pathway-level gene expression patterns across the WAP. Specifically, assembled contigs assigned to diatoms and annotated with KEGG Orthology identifiers (KO’s) that were annotated to a higher gene family or pathway level within a KEGG module (MO) were examined (Supplemental Fig. 4). Generally, diatom metabolic pathways were similar across the WAP, as these pathways are fundamental to cellular functioning in diatoms. Highly expressed genes included those associated with ribosome, central carbohydrate metabolism, carbon fixation, ATP synthesis, and spliceosome KEGG pathways. Yet, there were significant spatial variations in pathway metabolisms likely reflecting the changing oceanographic conditions in the region. Diatoms in coastal stations in southern regions appeared to invest proportionally more in these aforementioned pathways compared to those from slope and shelf regions in the North. In particular, *Actinocyclus* and other diatoms at station 100.040 invested heavily in carbohydrate metabolism, cysteine and methionine metabolism, protein processing and nucleotide biosynthesis metabolism (pyrimidine and purine). From the apparent high investment in genetic information processing, energy metabolism, and carbohydrate/lipid metabolism, we propose that these diatoms were experiencing high rates of translation and transcription, supported by C fixation and ATP synthesis to meet the high energy demands of rapidly blooming cells.

Although pathway level groupings of KOs provide information on broad metabolism, the number of KOs that have an associated MO annotation are typically small; in our study only 38% of KOs had a higher-level module annotation (i.e., 6% of all contigs). While still informative, this low level of annotation makes it difficult to draw definitive conclusions from module expression alone. Therefore, we queried the most variable KOs expressed across the WAP (transcripts with the highest log_2_-normalized variance across sites). Expression of the most variable diatom genes shifted along latitudinal and cross-shelf gradients (Supplemental Fig. 5). Coastal stations in the south had higher transcript levels for genes encoding nucleotide and energy production. Specifically, a pyrimidine precursor biosynthesis enzyme was highly abundant (THI5). Pyrimidines are particularly important in dividing cells as building blocks for nucleic acids, but they are equally important for many biochemical processes, including sugar (UDP) metabolism and vitamin B1 synthesis (58). Increased abundance of iron-containing photosynthesis subunits (PsbN and PsaC) as well as cytochrome c oxidase (COX) subunits, suggestive of increased ATP formation through cellular respiration, indicate conditions conducive to growth. In contrast, northern stations had more transcripts for stress related genes such as E3 ubiquitin-protein ligases and proteasome activator protein. Ubiquitination by E3 ligases regulates cell trafficking, DNA repair, and signaling, mediating the degradation of damaged proteins (59) which are processed by the proteasome. Transcripts for the gene encoding plastocyanin (PCYN), a protein used for Fe-independent transfer of electrons in photosynthesis, and a low Fe responsive aquaporin (AqpZ) (35) were also detected in the north, suggesting low Fe availability or other abiotic stressors. Slope stations demonstrated fewer transcripts compared to shelf or coastal regions.

To further contextualize spatial gene expression of diatoms within the WAP region, a weighted gene co-expression network analysis (WGCNA) was used to group KOs into clusters or modules (MEs) of highly correlated genes based on patterns of expression (49). These groups of similarly expressed genes were then correlated with the measured oceanographic parameters. Over 4000 genes detected in diatoms were clustered into nine MEs (Fig. 3A), reflecting correlations with oceanographic parameters along a coastal to slope gradient (Fig. 1B, C). MEs were assigned colors to distinguish them. The red and blue MEs, and to a lesser extent the yellow ME, contained highly expressed genes positively correlated with Chl *a*, primary production and phaeopigments (a degradation product of Chl *a*) and negatively correlated with temperature, salinity, DIC and macronutrient (i.e., nitrate/nitrite, and phosphate) concentrations. In contrast, the turquoise, green and brown MEs correlated with these parameters in the opposite direction, with highly expressed genes at stations where high nutrient concentrations and lower Chl *a* concentrations were present. Specifically examining the MEs with the strongest correlations in relation to Chl *a*, genes in the red module were uniquely highly expressed at southern coastal stations associated with the bloom of *Actinocyclus*, indicating that these blooming diatoms had a unique metabolic profile as examined through gene expression (Fig. 3B). In contrast, genes clustered in the turquoise ME showed increased expression in shelf and slope stations and were positively correlated with high salinity and nutrient concentrations, but showed opposing expression patterns in coastal stations where they were negatively correlated with these oceanographic parameters, highlighting the clear effects of the cross-shelf gradient in nutrients and salinity on gene expression (Fig. 3B).

**Figure 3.**
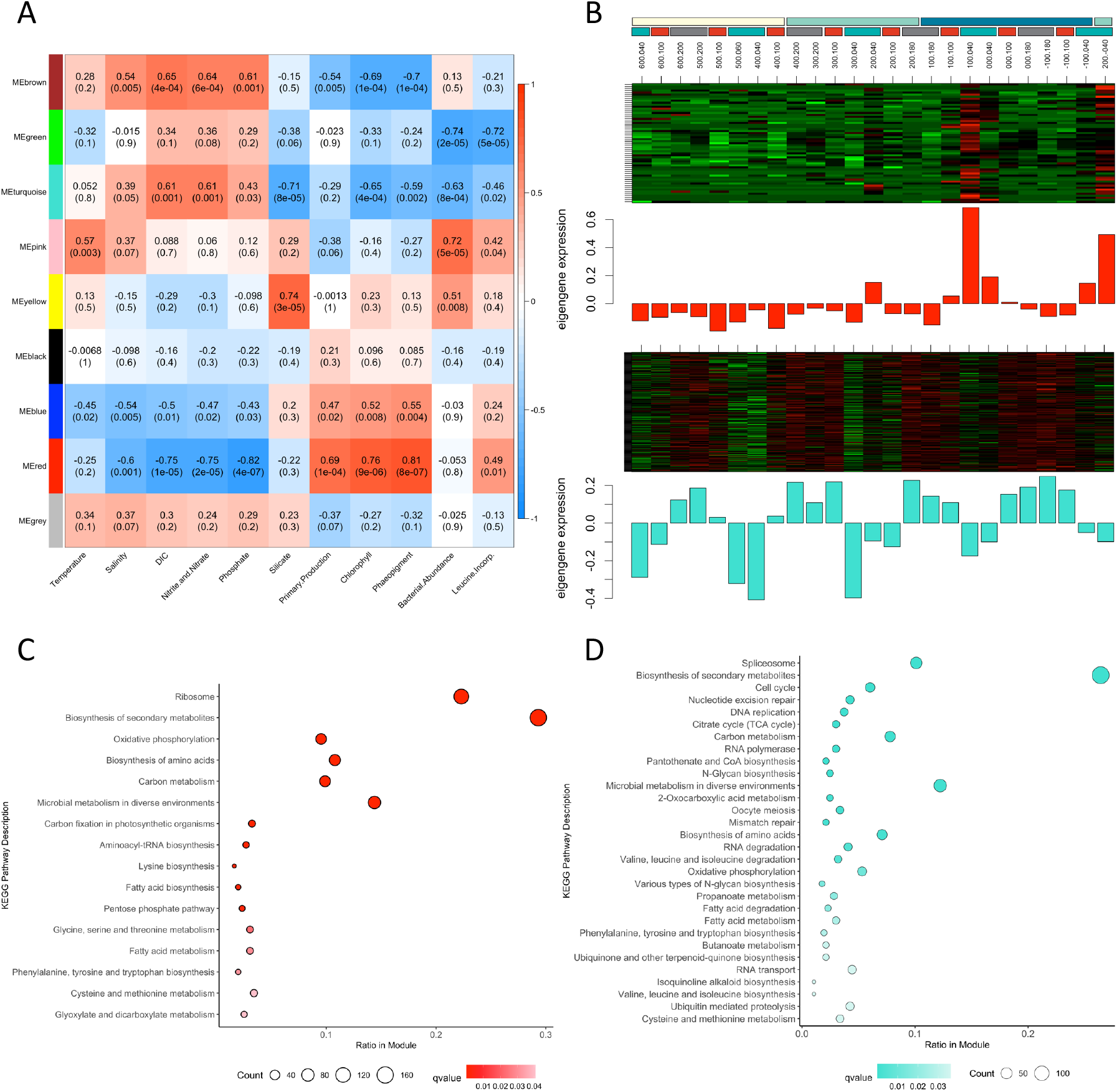
A) Weighted Gene Co-Expression Network Analysis (WGCNA) demonstrates the correlation between environmental parameters (x-axis) and groups of similarly expressed genes assigned to modules (ME) and designated by arbitrary colors (y-axis). Top numbers in each cell are Pearson’s correlation coefficients with bottom numbers representing p-values of the correlation test. Color of each cell indicates the correlation between ME and environmental parameters, with red indicating a positive correlation and blue indicating a negative correlation. Heatmap of transcripts per million (TPM) gene expression plotted alongside bar graphs of eigengene expression within B) the red module (ME) and C) the turquoise ME at each station. Dot plot of KEGG enrichment analysis of A) the red ME and B) the turquoise ME at the pathway level. Enrichment is based on the number of KO’s overrepresented in KEGG pathways compared to all annotated KO’s. Significance given to FDR adjusted p-values (qvalue) < 0.05, darker color indicate higher significance. Counts represent the number of KO’s detected in each pathway.

Each ME contained hundreds of genes associated with KEGG pathways and modules. By performing an enrichment analysis, distinct metabolic investments by diatoms in the red and turquoise ME’s, the two modules which displayed the most significant correlations with the oceanographic parameters measured, were determined (Fig. 3C-D). The *Actinocyclus* bloom stations, represented by the green ME, were significantly enriched in the KEGG Pathways: ribosome, carbon metabolism, C fixation, oxidative phosphorylation, and biosynthesis of secondary metabolites and amino acids. Similar patterns of enriched pathways have been observed in blooms of sea ice diatoms in the Wilkens Ice Shelf (45), dinoflagellate blooms in the Neuse River Estuary, NC (60), and Fe-induced diatom blooms in the NE Pacific Ocean (61), perhaps indicating a common response in blooming diatoms associated with growth and energy production. Although the cells at this station appear to have been blooming, the enrichment of glycine, serine, and threonine metabolisms indicate an effort to reduce photorespiration, or stress from reactive oxygen species (ROS), suggesting possible photoinhibition or perhaps the onset of bloom termination (62, 63).

In contrast, shelf and slope stations, represented by the blue turquoise ME, were enriched in KEGG pathways representing stress and regulation of transcription and translation: spliceosome, cell cycle, nucleotide excision repair, glycan biosynthesis, and DNA replication (Fig. 3D). The enrichment of glycan biosynthesis suggests increased formation of glycoprotein-rich extracellular matrices which may be important under freezing temperatures as has been observed in *F. cylindrus* (33). They are also part of a diverse family of transmembrane proteins that are widely implicated in allorecognition, including the establishment of symbiosis and microbial interactions (64). We speculate that increased expression of this pathway under nutrient limitation may foster interactions with beneficial bacteria or other microbes. In summary, these transcriptional patterns indicate blooming diatoms in southern coastal stations and potentially Fe-stressed diatom communities on the shelf and slope.

### Sea ice edge bloom of a large centric diatom

Analysis of remote sensing-derived surface Chl *a* concentration, cell densities, and gene expression revealed a bloom of the centric diatom *Actinocyclus* in the far south coastal regions of the Pal-LTER study area (Fig 2B, Fig. 4). Cell densities of *Actinocyclus* in surface waters peaked at station 100.040, achieving 2.2. x 10^4^ cells L^−1^. It is hypothesized that very large centric diatoms such as *Actinocyclus* can preferentially bloom in the Southern Ocean because their silica laden frustules provide ample defense against grazers such as krill, whereas smaller diatoms cannot build up high biomass due to higher mortality as a result of top-down controls (6). Blooms of large diatoms (> 20 µm) such as *Proboscia* and *Chaetoceros* have also been observed in pelagic and coastal regions of the Southern Ocean (20, 52, 65), particularly when Fe is replete and MLDs are shallow. Within the WAP region, both *Proboscia* and *Actinocyclus* blooms have been previously documented in Marguerite Bay (54, 66), associated with cold, fresher surface layers that follow from sea ice retreat. Furthermore, larger blooms occur following years with more sea ice and delayed retreat (15). Eventually, a wind-induced breakdown of the melt water layer causes a shift in the eukaryotic community from large blooming centric diatoms to smaller centric and pennate diatoms or cryptophytes (66).

**Figure 4.**
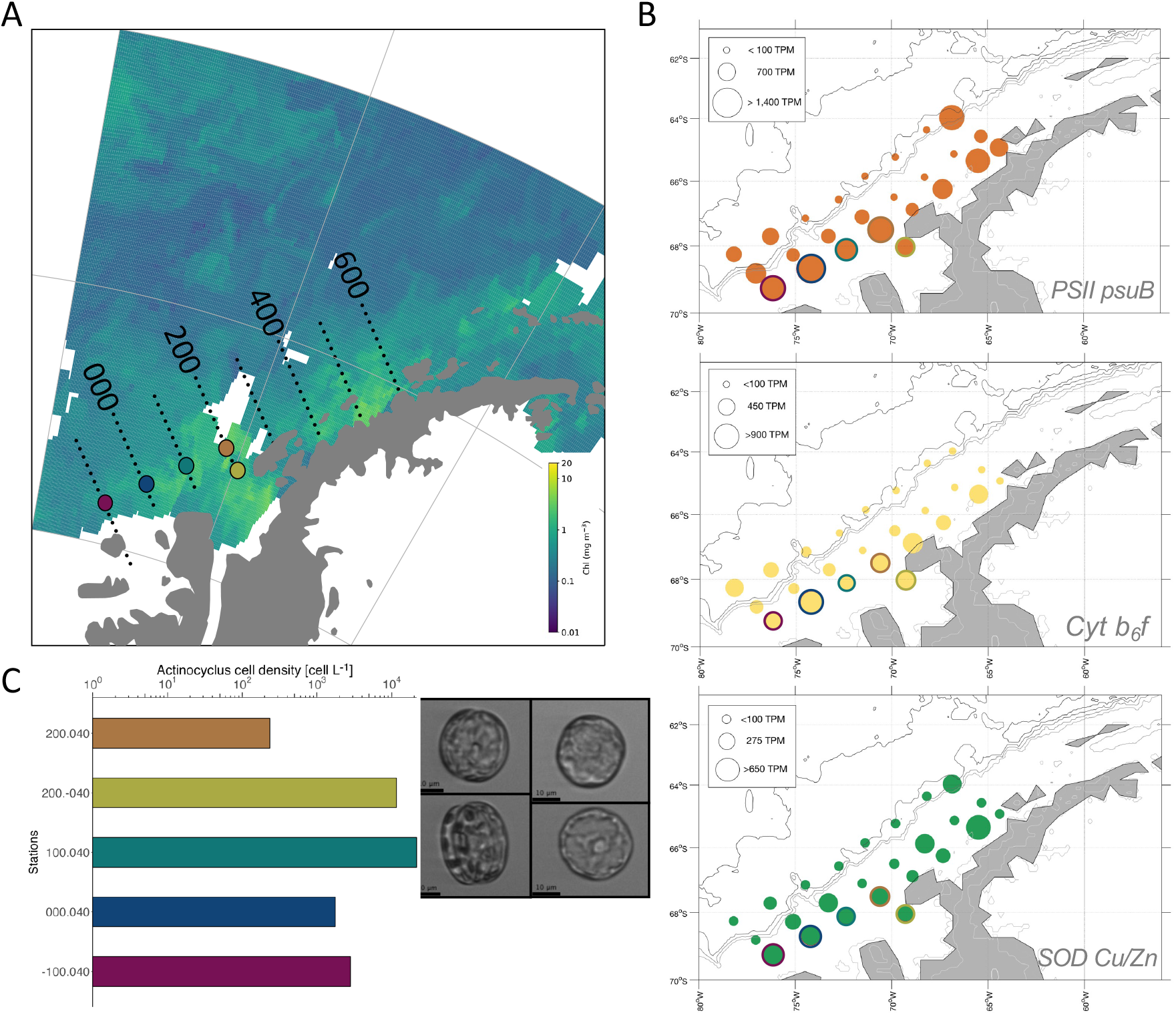
A) A large *Actinocyclus* sp. bloom in the Far South coastal region of the Pal-LTER grid, visualized with an 8-day average composite chlorophyll *a* satellite image (Jan 9-16, 2018). B) Log-transformed TPM of select *Actinocyclus* genes of interest of photosynthesis and ROS, that indicate active actively growing diatoms. Stations outlined in red indicate bloom locations. C) Cell counts and images of *Actinocyclus* sp. captured via an Imaging FlowCytobot from surface seawater samples.

Although *Actinocyclus* appears to be a frequent bloomer in the far south region, growth characteristics measured in laboratory isolates have demonstrated that members of this genus have relatively slower maximum growth rates compared to other polar centric and pennate diatoms (24, 67). *Actinocyclus* also appears to perform poorly under low iron conditions due to its high half-saturation constant for growth relative to iron concentration, and is susceptible to photoinhibition at high light levels (67, 68). A transcriptome analysis of the *Actinocyclus* gene repertoire suggests that it may lack many genes encoding proteins involved in low Fe coping strategies such as ferritin, an Fe-storage protein, or a complete high-affinity Fe uptake pathway (24). Thus, this diatom may be uniquely specialized for environments where cold meltwaters and elevated Fe levels are present, resulting in a boom and bust behavior that has been proposed as a life-style for some polar diatom species (65, 69) and is responsible for large amounts of C export (8).

Indeed, the *Actinocyclus* bloom we observed was characterized by the coldest and freshest waters measured along the WAP region in 2018 (Supplemental Fig.1). Satellite imagery of an 8-day composite during the sampling period demonstrate the high levels of Chl *a* in the region around station 100.040 (Fig. 4A), where maximum cell densities of *Actinocyclus* were observed among the bloom sites (Fig. 4B and 4C). The molecular investments made at the bloom stations were highest for ribosomal proteins, oxidative phosphorylation and photosynthesis (Supplemental Fig. 5, 6). The region appeared to be naturally Fe replete as evidenced by the high expression of genes for Fe-requiring proteins, such as photosystem II subunits and cytochrome b6f, and relatively low expression of iron-starved induced proteins (ISIPs). Diatoms are known to have an increased ability to capitalize on newly available nutrient resources, with high initial investment in energy metabolism (70). It may be that while *Actinocyclus* exhibits slower growth rates and has more specialized niche requirements than diatoms such as *Fragilariopsis* or *Pseudo-nitzschia*, it can quickly allocate resources to growth. However, its large size, as demonstrated through images captured on an Imaging Flow Cytobot (Fig. 4C) and high silica requirement (58) indicate its main strategy is likely predator avoidance in a specialized niche. Interestingly, an *Actinocyclus* peroxiredoxin, an FK506-binding protein (FKBP) and a cold-shock protein (CspA) were highly expressed at non-bloom, shelf and slope stations. FKBPs are a family of conserved proteins involved in diverse cellular functions including protein folding, cellular signaling, apoptosis and transcription (71) whereas CspAs in bacteria such as *E. coli* function as an RNA chaperone at low temperatures (72).

### Evidence for iron limitation in WAP diatoms

While light availability and degree of water column stratification are central to regulating productivity in coastal waters, micronutrients, especially Fe, have been suggested to be particularly critical in the Southern Ocean and offshore waters of the WAP region (4, 23, 37). Several lines of evidence for Fe limitation in the WAP region exist. First, a cross-shelf gradient of dissolved and particulate iron has been observed, with particularly low concentrations (< 0.1 nmol kg^−1^) widespread in shelf and slope stations, especially in the northern half of the sampling grid (23). Second, severe Fe limitation has been strongly suggested through the measurement of phytoplankton photophysiology, with decreased photosynthetic energy conversion efficiencies and increased decoupled light harvesting chlorophyll-protein antenna complexes in shelf and slope waters, signatures typical of Fe limitation (37, 73). And third, upwelled UCDW waters in offshore and slope areas result in subsurface phytoplankton maxima (18) in contrast to the surface blooms that dominate nearshore waters. Thus, coastal phytoplankton benefit from waters with entrained Fe from sediments, sea ice, or from sources of meteoric water (glacial melt and precipitation) (23). However, until now, molecular evidence of Fe limitation from ‘omic approaches is generally lacking in this region.

To better understand if polar diatoms experienced a cross shelf gradient in Fe availability, we examined the expression of genes encoding Fe-responsive proteins such as those for Fe-dependent proteins and their functional replacements, as well as those proteins involved in ROS mitigation and nutrient acquisition. We focused on two dominant diatom genera, as measured through taxonomically annotated transcripts: *Fragilariopsis* and *Pseudo-nitzschia*, both of which are also known to respond to Fe-enrichment in HNLC regions around the globe. Unlike *Actinocyclus*, these two diatoms were detected at all sampled stations (Fig. 2B). Expression of Fe-related genes reflected an on/offshore gradient with most coastal sites (Fig. 5A, B). Relatively high expression of the genes encoding the iron starvation induced protein 1 (ISIP1) suggests iron limitation occurs in shelf waters, regardless of latitudinal location along the WAP. ISIP1 is speculated to assist diatoms in taking up hydroxamate siderophores via endocytosis (74) and transcripts for the protein are more common in Southern Ocean diatoms (24) compared to temperate diatoms. In *F. kerguelensis*, ISIP1 was highly expressed under Fe limitation but was also expressed under low light conditions (35). This may explain its constitutive expression, even at low levels, across all stations in the WAP region. While phytotransferrin (previously ISIP2a) has also been proposed as a molecular indicator for Fe availability in polar diatoms (75), expression was only detected in *Pseudo-nitzschia* at a few stations. It may be that ISIP1 could serve as a more useful molecular indicator of Fe limitation in this region.

**Figure 5.**
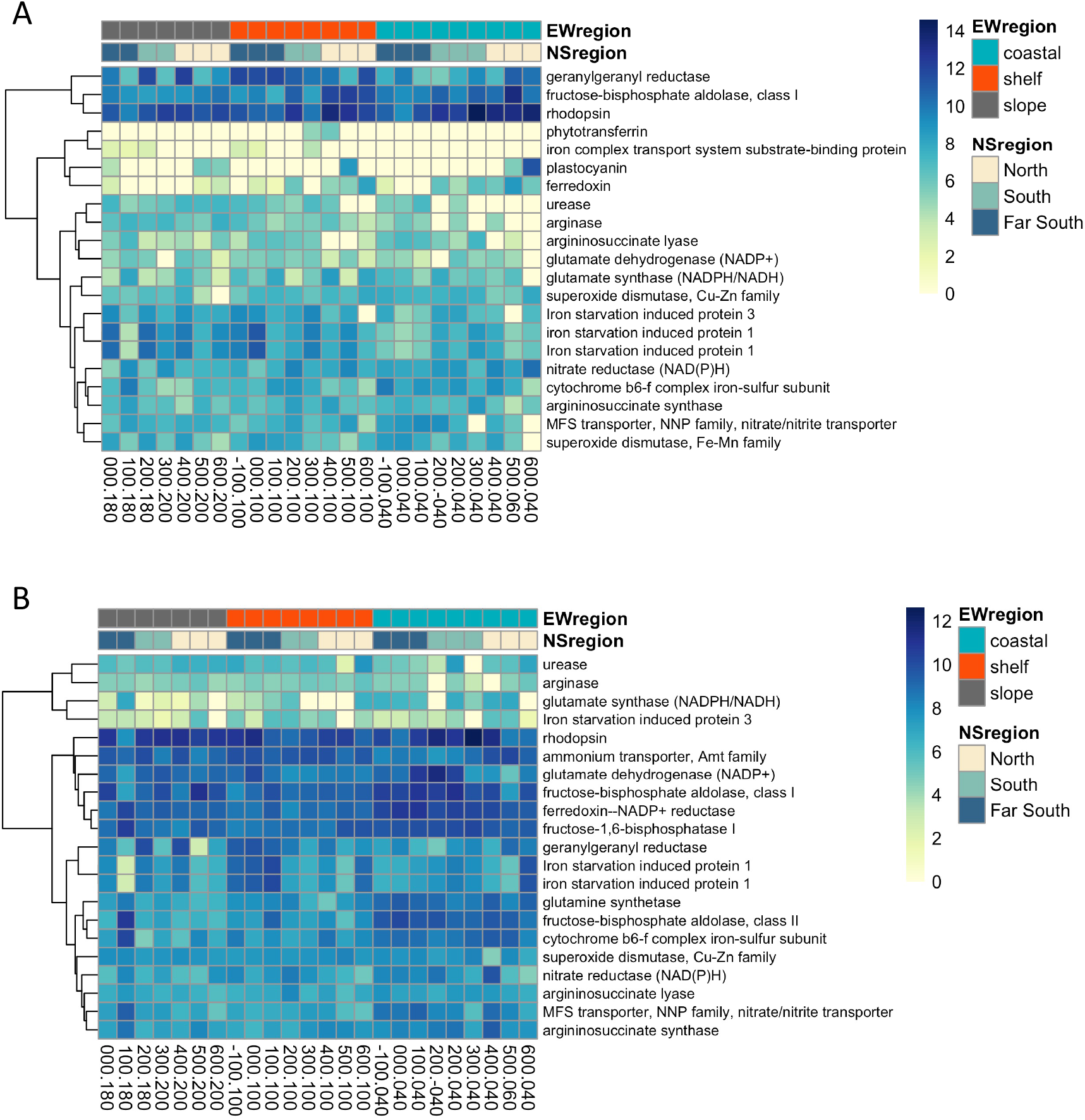
Heatmap of transcripts recruiting to KO’s relating to N uptake, Fe homeostasis, and photosynthesis in the diatoms A) *Pseudo-nitzschia* and B) *Fragilariopsis*. Scale indicates transcripts per million. Dendrograms show similarity in transcript abundances determined with Euclidean distances and hierarchical clustering.

Stations that had increased ISIP1 expression also had higher abundances of a geranylgeranyl reductase (ChlP), a key precursor and regulator of Chl *a* synthesis and tocopherol production (62). ChlP may be useful in remodeling the photosynthetic apparatus to cope with Fe limitation at offshore stations where Chl *a* concentrations were reduced. As tocopherol is a strong antioxidant (62), ChlP is also likely involved in the removal of ROS that stems from the inefficient activity of an Fe-limited photosynthetic electron transport chain. Interestingly, RHO expression was also abundant everywhere along the WAP region, but especially at coastal sites in the northern half of the grid, and often coincided with reduced ISIP1 expression. RHO may not be driven strongly by Fe alone in this region (32), but rather by high light levels in shallow MLDs of coastal stations. *Actinocyclus* RHO also followed this same pattern (Supplemental Fig. 8). Other photosynthesis Fe requiring protein-encoding genes were highly expressed at coastal stations in both diatoms, including ferredoxin, cytochrome b6f and a vitamin B1 synthesis gene THIC (Fig. 5).

The expression of genes encoding nutrient acquisition and metabolism were queried along the WAP region. Nitrogen acquisition and assimilation requires Fe and energy and should show contrasting patterns along a cross-shelf iron gradient. Both nitrate reductase (NR) and a nitrate/nitrite transporter (NRT) were more highly expressed in coastal waters mirroring the expression of ISIP1 in shelf waters. The ornithine-urea cycle is important in recycling nitrogen throughout the cell (62), yet the genes we detected were not particularly abundant throughout the WAP region, the exception was urease which had relatively higher expression in shelf and slope stations waters in both diatoms. Urease may be helpful in transporting urea or other reduced N compounds as it does not have an Fe requirement. A urea transporter and urease were additionally detected in a metatranscriptome from the north Bransfield Strait (44). Taken together, these molecular data suggest that diatoms in coastal regions are experiencing nutrient replete conditions with high irradiances, and within a relatively short distance from shore (~100 km) diatoms and other phytoplankton groups may be experiencing Fe limitation in an HNLC-like environment.

### Conclusions

Metatranscriptomes analyzed along the Pal-LTER survey elucidated the molecular physiology and metabolic flexibility of diatoms along the diverse habitats of the Western Antarctic Peninsula. Distinct phytoplankton communities and metabolisms were observed and closely mirrored the strong gradients in oceanographic parameters that existed from coastal to offshore regions. Diatoms were abundant in coastal regions especially in the Far South, where cold and fresher waters were conducive to a diatom bloom. Under these favorable conditions, the large centric diatom *Actinocyclus* was investing heavily in growth, energy production, carbohydrate and amino acid and nucleotide biosynthesis pathways, resulting in a uniquely expressed metabolic profile in this Far South region.

In waters of the shelf and slope regions, we observed strong molecular physiological evidence for Fe limitation, supporting previous chemical and physiological studies. Diatoms in these regions employ ISIPs, a geranylgeranyl reductase, an aquaporin, and urease, among other strategies, while limiting the use of Fe-containing proteins, as inferred through gene expression analysis. The findings here reveal functional differences in diatom communities and is a first step in providing conclusive evidence for iron limitation and how certain ecologically dominant members of the phytoplankton community cope under these conditions.

The ecological success of diatoms along the WAP is likely related to their unique physiological adaptations, their resistance to a co-limitation of iron and light, and their unique molecular adaptations. The gene expression patterns observed here demonstrate the strong influence oceanographic factors have in shaping disparate metabolic profiles of diatoms along the peninsula. This spatial analysis of phytoplankton composition and diatom metabolism in this important region of the Southern Ocean will contribute to our understanding of how these critical marine ecosystems might shift in future climate change scenarios, especially along a latitudinal gradient of the WAP, and will serve as a critical baseline for future studies of molecular physiology of natural plankton assemblages.

## Materials and Methods

### The study region

The Palmer Long Term Ecological Research (Pal-LTER) study region is located along the western coast of the Antarctic Peninsula, encompassing an area of approximately 140,000 km^2^ (Fig. 1A). Individual stations are arranged in a grid from the coast to offshore, running perpendicular to the peninsula, with major grid lines spaced 100 km apart and individual stations spaced 20 km apart along each line (15, 45). The region is divided further based on latitudinal, hydrographic and sea ice conditions (46). The Northern region consists of lines 400 – 600, the South consists of lines 200 and 300, and the Far South consists of lines −100 – 100. Following an onshore to offshore gradient, stations .000 – .040 are within the coastal region, station .100 is within the shelf region, and stations .180 and .200 are over the continental slope. Samples were collected during the Pal-LTER 2018 austral summer research cruise (30 December 2017 – 12 February 2018) aboard the ARSV *Laurence M. Gould*.

### Oceanographic data and sampling

All oceanographic data are publicly available through the Pal-LTER DataZoo repository at http://pal.lternet.edu/data, with detailed descriptions of sample collection and processing methods. In addition to sea surface temperature and salinity, chlorophyll *a* (Chl *a*), primary production (PP), bacterial production and abundance, and inorganic nutrients were used in this analysis (Supplemental Table 1). Cell densities of the centric diatom *Actinocyclus* were estimated from discrete whole seawater samples passed through an Imaging FlowCytobot (47). For RNA, approximately 2-4 L of surface seawater was collected (depending on phytoplankton biomass) and filtered through a 47 mm 0.2 µm Supor filter (Millipore). Each filter was preserved in 1 mL of RNA-later and stored at −80°C. Samples were kept frozen on dry ice during shipment to the University of North Carolina at Chapel Hill. Prior to RNA extraction, filters were thawed and cut in half with one half of the filter used for RNA sequencing analysis while the other was archived.

### RNA extraction, sequence library preparation and sequencing

Total RNA was extracted with the RNAqeuous 4PCR Kit (Ambion) according to manufacturer’s instructions with the addition of an initial one-minute bead beating step using acid-washed sterile 425-600um glass beads (Sigma Aldrich) to ensure cells were mechanically removed and disrupted from the filter. Samples were eluted in 40 μL of sterile H_2_O and stored in −80°C. Residual genomic DNA was removed by incubating RNA with deoxyribonuclease (DNase) I at 37°C for 45 minutes and purified by DNase I inactivation reagent (Life Technologies). Some samples required multiple incubations with DNase I. Sample concentrations and RNA integrity numbers (RIN) were determined using an Agilent Bioanalyzer 2100. RIN values were between 3.9 and 7.4. Messenger RNA (mRNA) libraries were generated with *ca*. 2 μg of total RNA, using a poly-A selection primarily selecting mRNA of the eukaryotic plankton community, and prepared with the Illumina TruSeq mRNA Library Preparation Kit. Samples were individually barcoded and pooled prior to sequencing on a single lane of the Illumina HiSeq 4000 platform at Genewiz Sequencing Facility (S. Plainfield, NJ). Sequencing resulted in *ca*. 15 million 2×150 bp paired-end reads per sample (Supplemental Table 2). Sequences were submitted to the NCBI Sequence Read Archive under accession number PRJNA877830.

### Metatranscriptome assembly and annotation

Sequences were quality filtered with Trimmomatic v0.38 and summary statistics were generated before quality filtering with FastQC v0.11.8, and after Trimmomatic with MultiQC v1.9. Trimmomatic (paired-end mode) removed Illumina adapters and used a 4 bp sliding window to remove quality scores below 20 and to keep sequences longer than 36 bp. Paired-end sequences from each sample that passed quality control were individually assembled using rnaSpades v3.14.1 resulting in 25 individual assemblies of contigs. The resulting contigs from all samples were combined into one consensus assembly using cd-hit-est v4.8.1 by clustering all samples at 100% identity, with a short word filter of size 11 (k-mer), and an alignment coverage of 98% for the shorter sequence (-aS). This consensus assembly was representative of all expressed genes of the microeukaryote community along the WAP and was used in the subsequent annotation and read count steps.

Taxonomic annotations of consensus sequences were assigned using Diamond BLASTX v2.0.4 based on homology (e-value cutoff of 10^−5^) to PhyloDB v1.075, a custom database curated by the Allen Lab (Scripps Oceanographic Institute), consisting of eukaryotic genomes and the transcriptomes from the Marine Microbial Eukaryote Transcriptome Sequencing Project (MMETSP) (48). We further supplemented the database with eight polar transcriptomes isolated from the WAP and previously sequenced (24). For this study, the individual polar transcriptomes were translated to protein space with GeneMarker-ES v4.61 and functionally annotated with EggNOG-Mapper v2.

For functional annotations, consensus sequences were searched against the Kyoto Encyclopedia of Genes and Genomes (KEGG) database v2018 by similarly using Diamond BLASTX and an 10^−5^ e-value. Annotation of proteins based on KEGG database classifications https://github.com/ctberthiaume/keggannot was performed using the python package Keggannot. Best hits to a functional protein and assigned KEGG Orthology (KO) identifiers and their higher-level classification, e.g. modules (MO) and Class 3 categories were obtained. Because KEGG only provides KOs for functionally annotated genes, KEGG gene definitions for certain targeted genes (e.g., iron-starved induced proteins (ISIPs) and rhodopsin sequences) were manually performed.

Trimmed reads from individual assemblies were aligned to the consensus assembly with Salmon v0.14.0 to obtain read counts. Read counts for consensus sequences that had taxonomic identifications from PhyloDB were used to evaluate plankton community composition of annotated transcripts. Read counts of consensus sequences that had an assigned KO identification number were used for functional annotations at the KEGG gene, module, and Class 3 level. Read counts of contigs sharing identical taxonomic and functional assignments were summed together. To account for shifts in community composition, reads were normalized within groups at the level of taxonomic interest (e.g., dinoflagellates, diatoms, *Pseudo-nitzschia*, etc.) using transcripts per million (TPM).

### Statistical analyses and data visualization

Principle coordinate analysis (PCA) was used with oceanographic data and specific genes of interest to visualize the spatial variability of samples along the WAP with the R package factoextra. To test if regionally grouped samples were statistically different, a permutational multivariate analysis of variance (PERMANOVA) test was conducted. A Canonical Correspondence Analysis (CCA) was used to determine the variance contribution of oceanographic factors on taxonomically annotated diatom RNA-sequences in the R package vegan. Raw counts were log transformed to reduce weight given to taxa with small counts, and oceanographic variables that were strongly correlated together were removed from analysis leaving the following to be included in the CCA: sea surface temperature, sea surface salinity, dissolved inorganic nitrogen (nitrate and nitrite), phosphate, silicic acid, chlorophyll a concentrations, primary production and bacterial production. A PERMANOVA was used to determine significance of the model. To visualize TPM gene expression at the KO, MO and Kegg Class 3 levels, heatmaps were produced with the R package pheatmap and clustered using Euclidean distance and hierarchical clustering. Finally, to identify groups of highly correlated genes that co-occur across the WAP region and to quantify effects of oceanographic variables on those groups, we used a Weighted Gene Correlation Network Analysis (WGCNA) (49). WGCNA clustered KOs, and their associated log_2_ TPM-normalized counts, into eigengene modules (ME, designated by arbitrary colors), representing the first principal component of the module matrix of similarly expressed genes. Using a minimum module size of 30 and dynamic tree cutting, 10 MEs emerged, two of which were combined into the ‘grey’ ME. Finally, MEs were correlated with oceanographic and biochemical data by pairwise Pearson correlation coefficients and corresponding p-values. A KEGG Class 3 enrichment test was performed with the R program clusterProfiler.

## Acknowledgments

We are grateful to the crew of the ARSV Laurence M. Gould, scientific support personnel, and Palmer LTER staff for their assistance in the field and lab. This research was funded by the National Science Foundation Grants OPP1745036 (to A.M.) and PLR1440435 (to O.S.). C.M.M. was primarily supported by a Gates Millennium Fellowship. We thank Yajuan Lin and Sarah Davies for insight on sequence analysis and helpful comments on the manuscript.

