## Supplemental figures and tables for "Molecular physiology of Antarctic diatom natural assemblages reveals multiple strategies contributing to their ecological success"

*Corresponding author: Adrian Marchetti

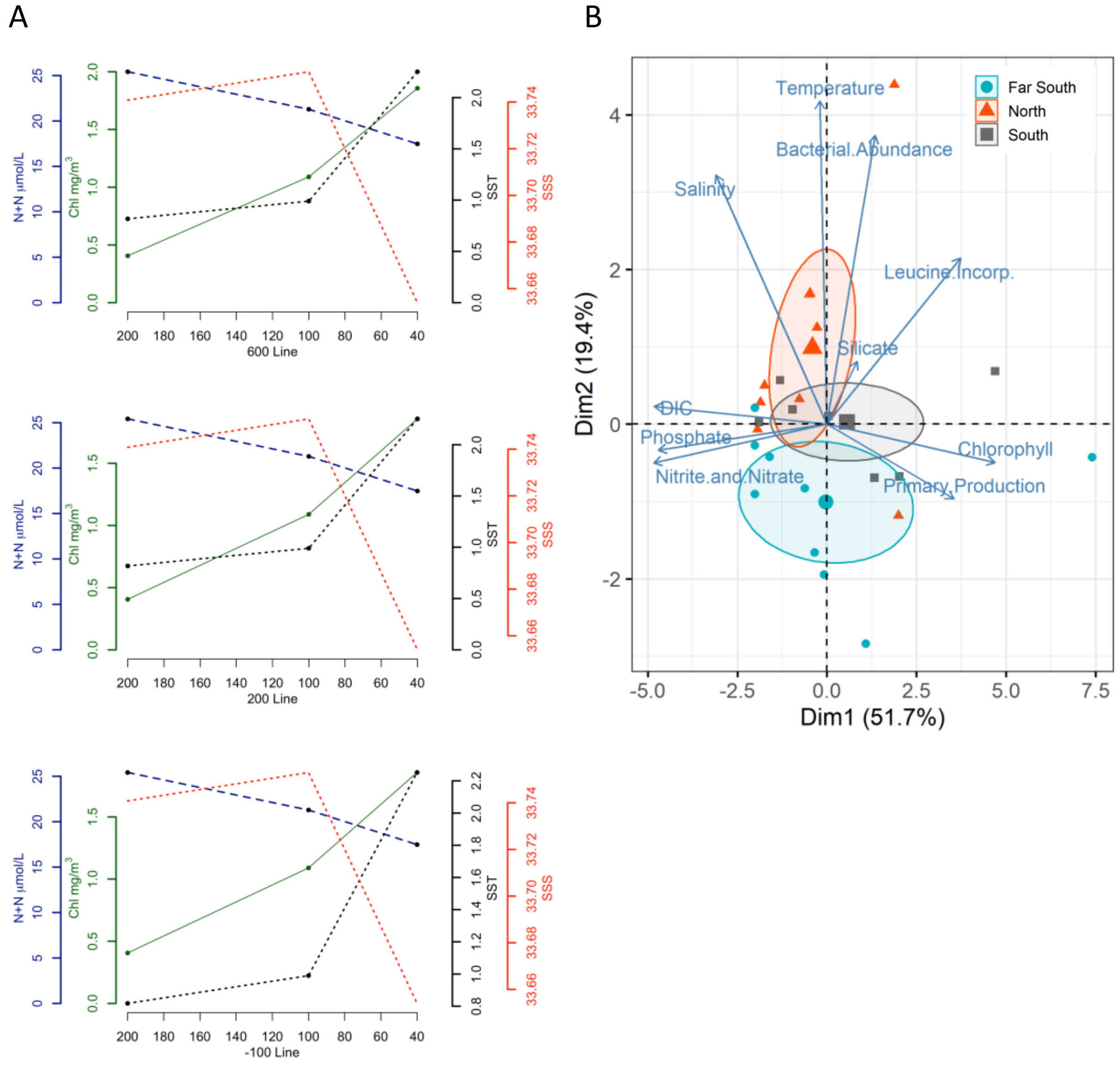


**Fig. S1.** Supplemental Figure 1. A) Oceanographic parameters on lines 600, 200 and -100 of the WAP. B) PCA bi-plot of environmental measurements with clustering of stations according to a latitudinal gradient from North to Far South regions

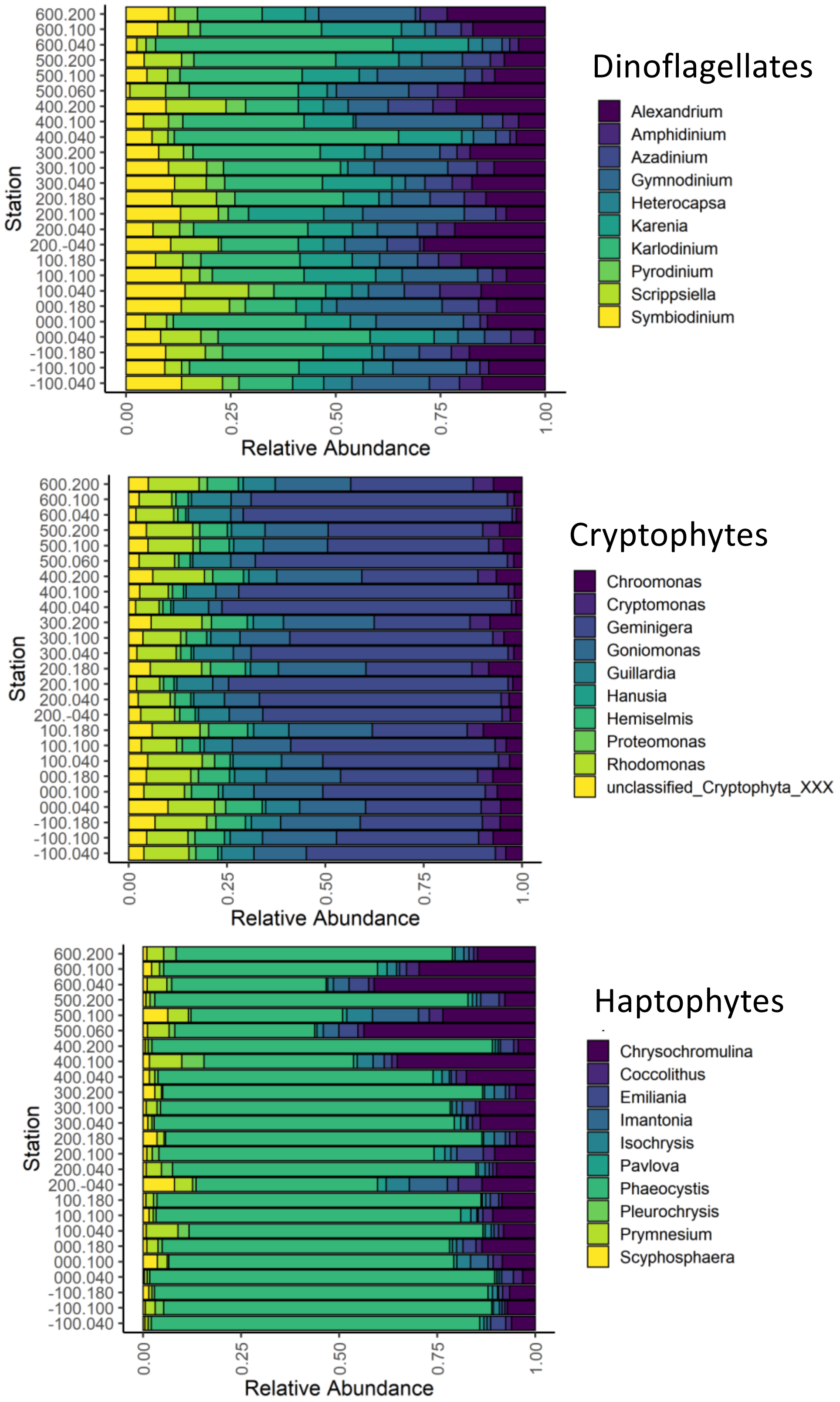


**Fig. S2.** Relative abundance of the composition of each major subdivision of plankton. Species included were present in at least 1% of all samples.

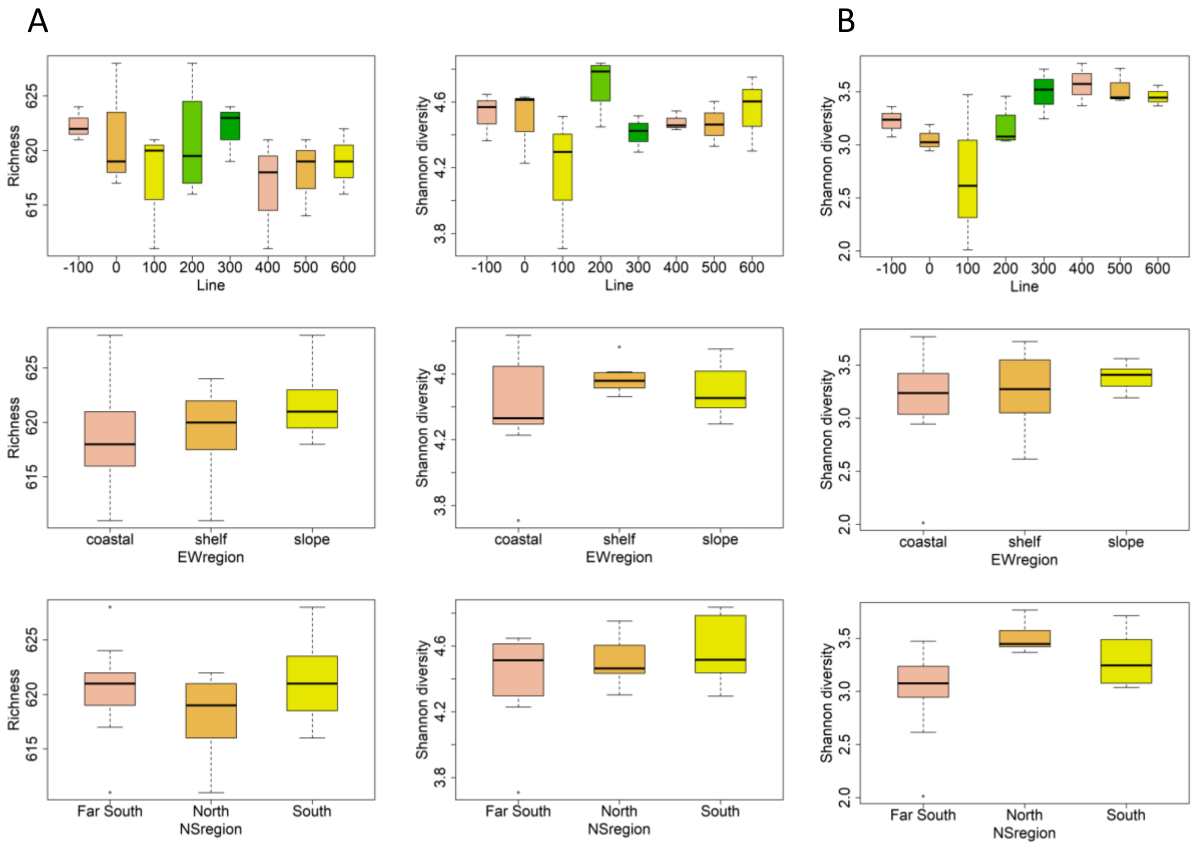


**Fig. S3.** A) Richness and Shannon-Weiner diversity of all eukaryotic phytoplankton across different regions. B) Diatom diversity as a function of different regions along the WAP. Richness not included as equivalent species were found in all regions.

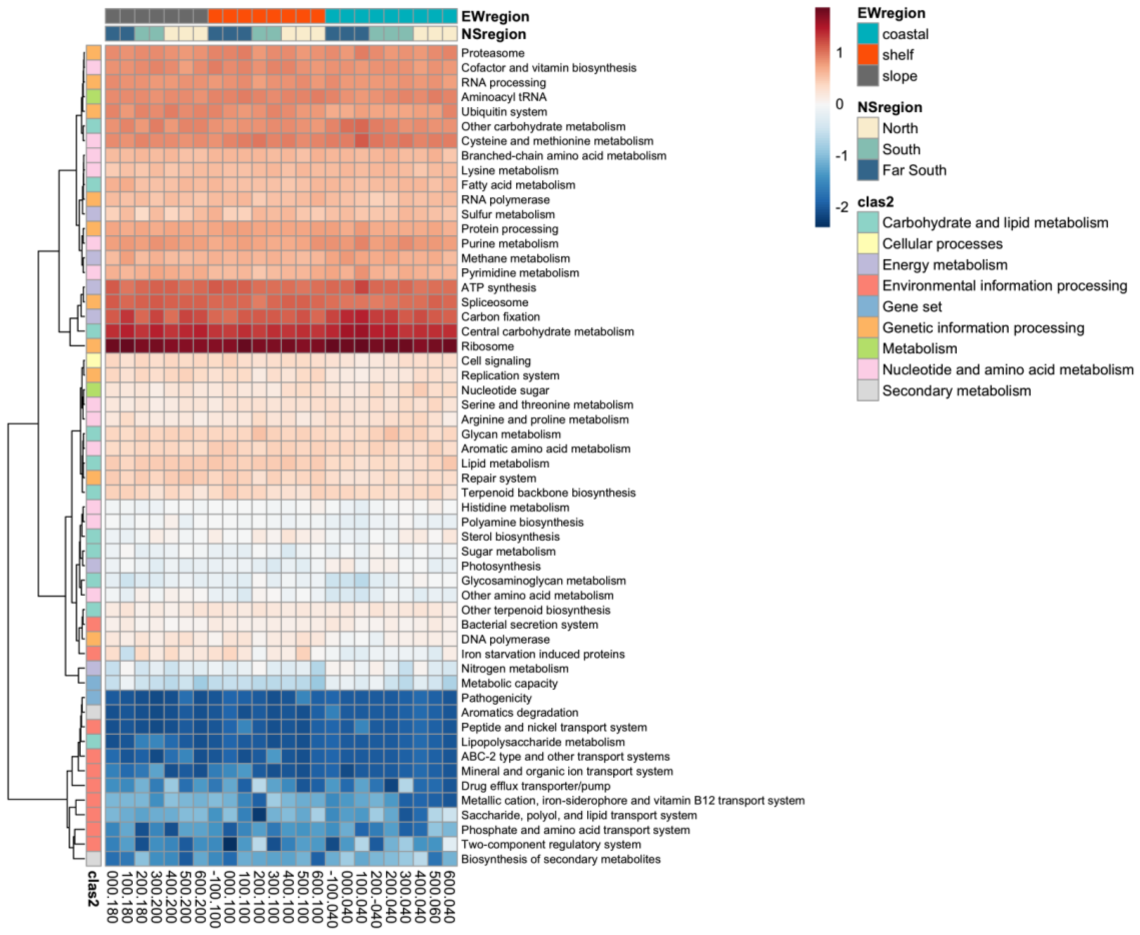


**Fig. S4.** Heatmap of transcript proportions recruiting to KEGG Class 3 broad scale categories (rows) for all diatom groups. Scale indicates high (red) or low (blue) transcript deviations from the mean (z-score) calculated across samples (columns). Dendrograms show similarity in transcript abundances determined with Euclidean distances and hierarchical clustering. Iron starvation induced proteins (pTF, ISIP1, ISIP2b, ISIP3) manually added as their own module.

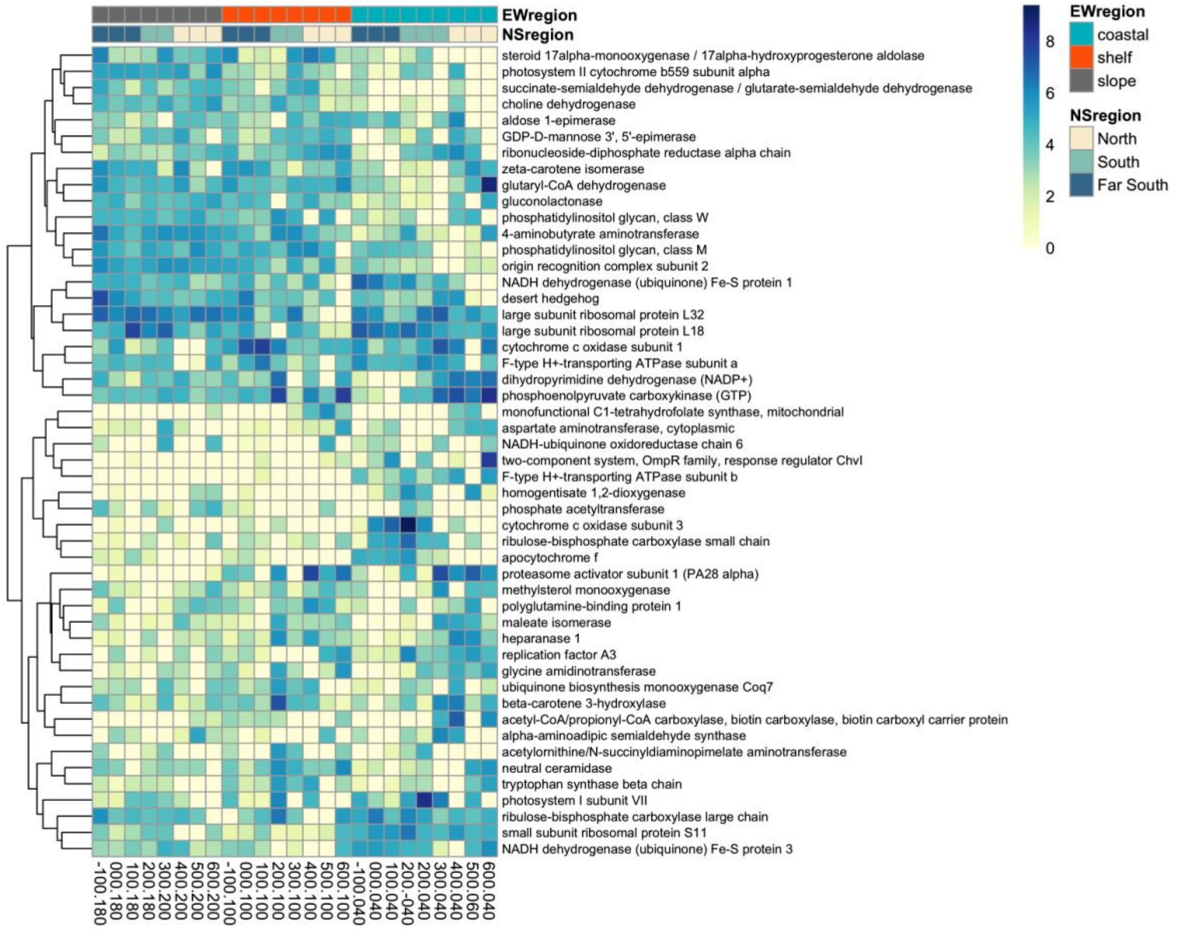


**Fig. S5.** Heatmap of diatom specific transcripts recruiting to the top 50 most variable KOs (rows) along the WAP. Scale indicates transcripts per million. Dendrograms show similarity in transcript abundances determined with Euclidean distances and hierarchical clustering

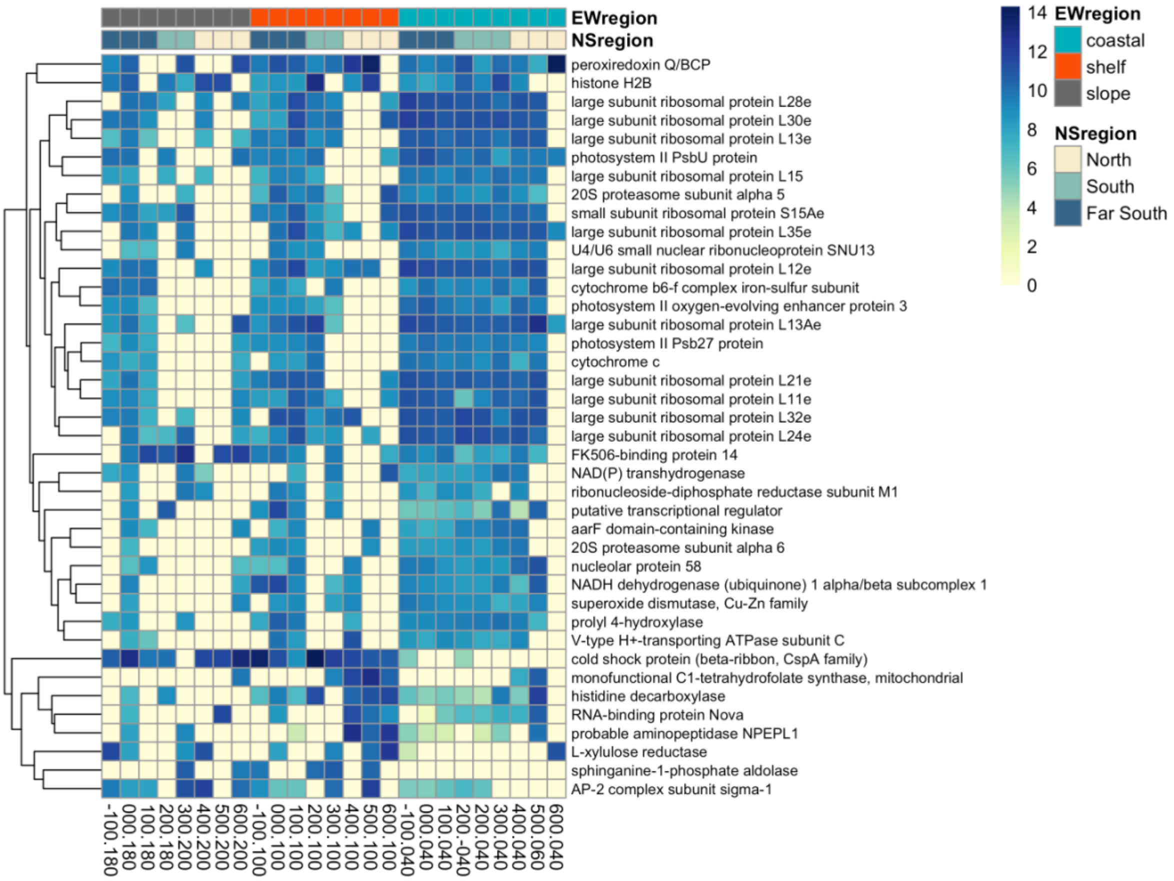


**Fig. S6.** Heatmap of transcript proportions recruiting to the top 50 most variable KOs (rows) for *Actinocyclus* sp. Scale indicates transcripts per million. Dendrograms show similarity in transcript abundances determined with Euclidean distances and hierarchical clustering. Stations highlighted in red indicate *Actinocyclus* sp. bloom stations.

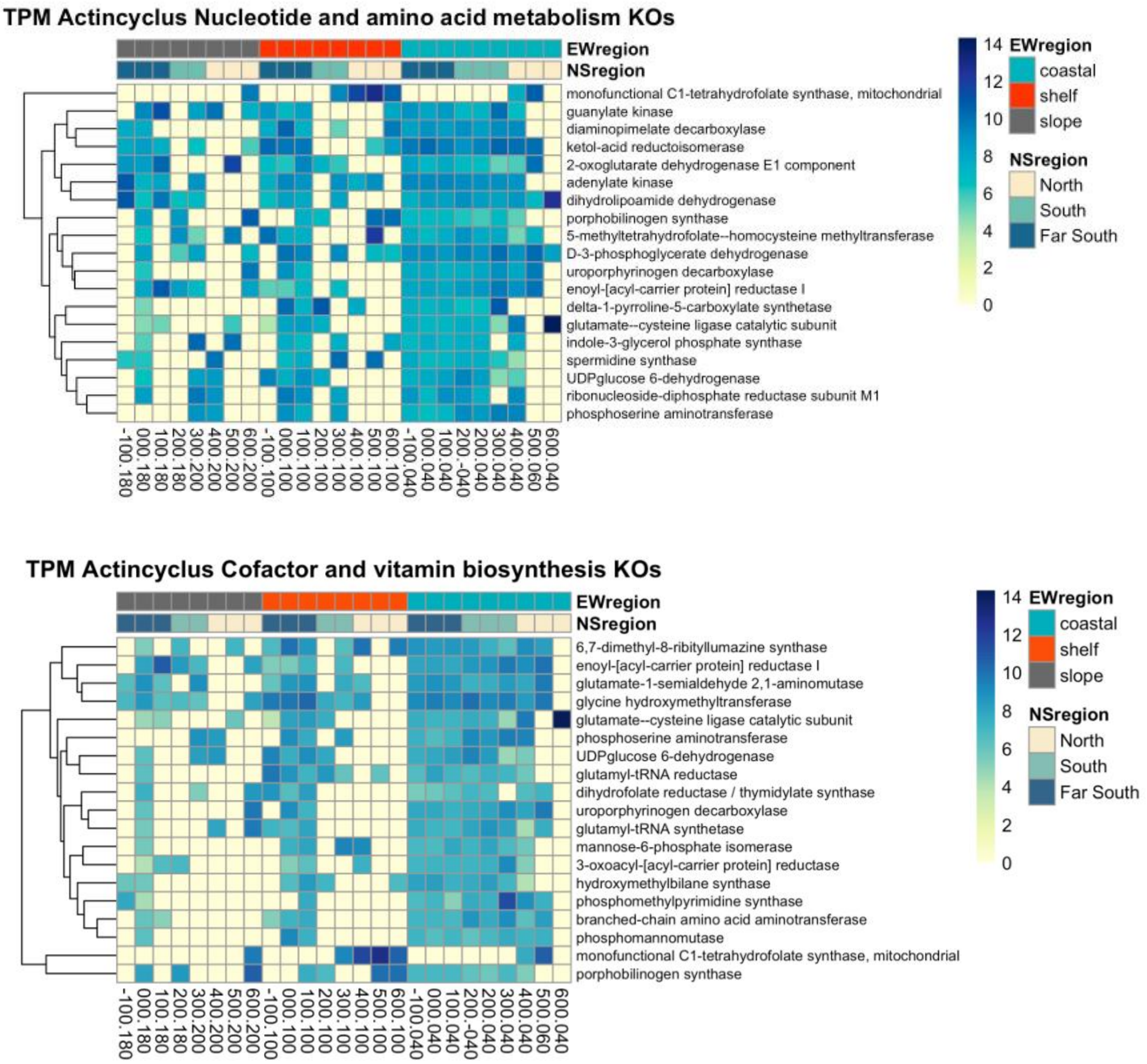


**Fig. S7.** *Actinocyclus* metabolism specific gene expression.

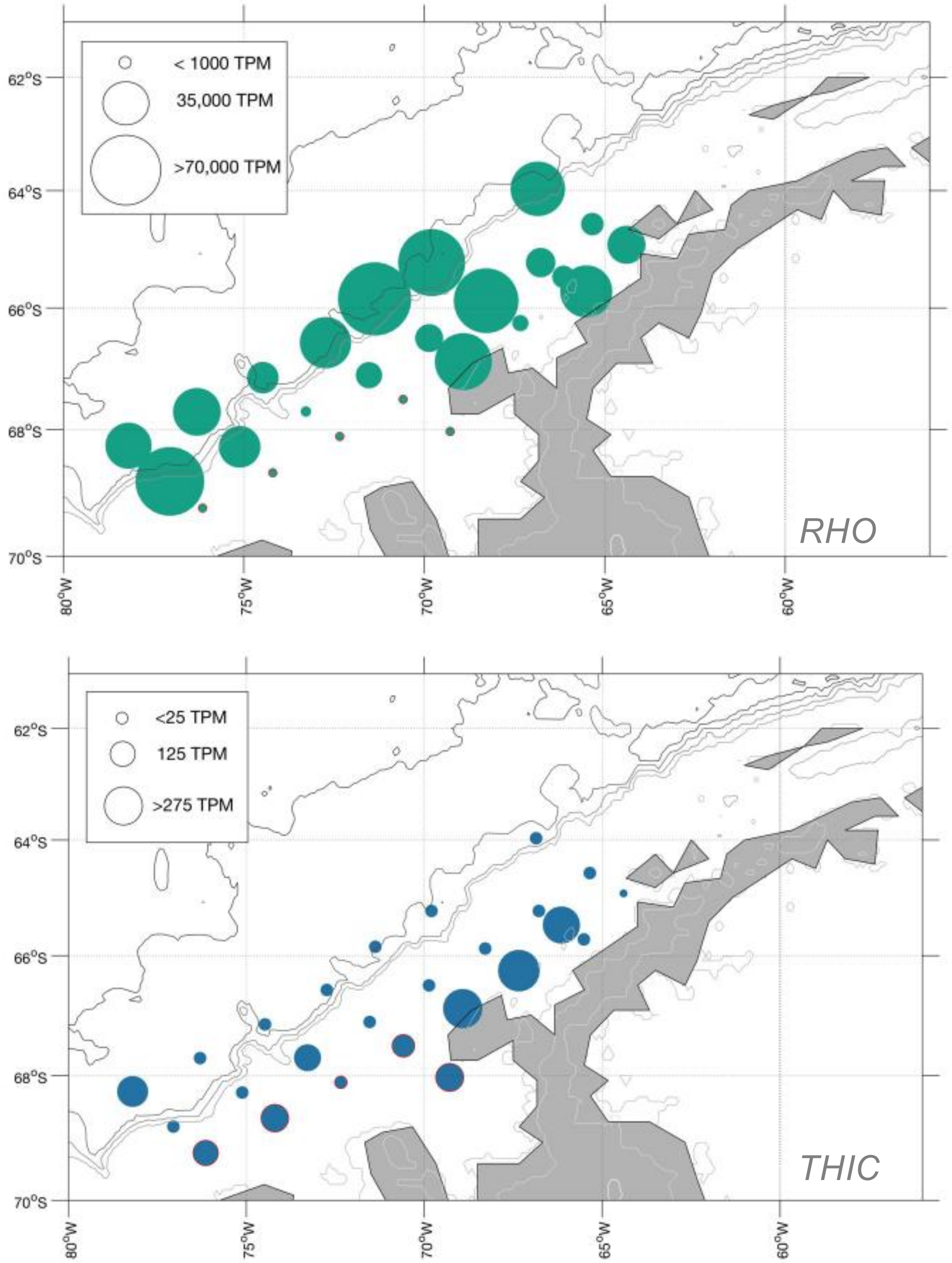


**Fig. S8.** Log-transformed TPM of select *Actinocyclus* RHO and THIC genes that indicate active actively growing diatoms. Stations outlined in red indicate bloom locations

**Table S1.** Oceanographic parameters measured during the 2018 Pal-LTER. North-South (NS) regions and East-West (EW) regions are listed for each station according to Steinburg et al. 2013.

| Station | Latitude | Longitude | NS region | EW region | °C | Salinity | DIC (umol kg$^{-1}$) | Nitrite/Nitrate (µmol L$^{-1}$) | Phosphate (µmol L$^{-1}$) | Silicate (µmol L$^{-1}$) | Primary Production (mg C m$^{-2}$ d$^{-1}$) | Chlorophyll *a* (µg L$^{-1}$) | Bacterial Abundance (cell L$^{-1}$) | Bacterial Production (pmol L$^{-1}$ h$^{-1}$) | Phaeopigment (µg L?1) |
|---|---|---|---|---|---|---|---|---|---|---|---|---|---|---|---|
| 600.040 | -64.93321 | -64.40059 | North | coastal | 2.25 | 33.65 | 2109 | 17.48 | 1.30 | 55.26 | 6.55 | 1.86 | 2.71E+09 | 97.73 | 0.4 |
| 600.100 | -64.57501 | -65.33957 | North | shelf | 0.99 | 33.75 | 2146 | 21.29 | 1.58 | 55.04 | 54.04 | 1.09 | 1.86E+09 | 8.30 | 0.3 |
| 600.200 | -63.9653 | -66.85524 | North | slope | 0.82 | 33.74 | 2157 | 25.42 | 1.68 | 41.96 | 25.55 | 0.41 | 5.19E+08 | 9.59 | 0.11 |
| 500.200 | -64.61099 | -68.29379 | North | slope | 0.52 | 33.74 | 2157 | 24.53 | 1.66 | 38.58 | 8.37 | 0.53 | 3.61E+08 | 7.06 | 0.15 |
| 500.100 | -65.23418 | -66.77544 | North | shelf | 0.87 | 33.70 | 2144 | 21.62 | 1.58 | 55.71 | 14.85 | 1.16 | 1.58E+09 | 33.09 | 0.44 |
| 500.060 | -65.47801 | -66.15043 | North | coastal | 0.98 | 33.59 | 2135 | 19.78 | 1.57 | 57.70 | 29.76 | 1.21 | 1.62E+09 | 5.05 | 0.38 |
| 400.040 | -66.25231 | -67.33584 | North | coastal | 0.03 | 33.19 | 2098 | 18.43 | 1.45 | 57.87 | 30.95 | 4.01 | | 51.47 | 1.31 |
| 400.100 | -65.87719 | -68.28142 | North | shelf | 0.54 | 33.77 | 2161 | 22.23 | 1.61 | 55.73 | 101.59 | 0.32 | 6.79E+08 | 16.97 | 0.09 |
| 400.200 | -65.23804 | -69.79766 | North | slope | 1.35 | 33.71 | 2150 | 23.58 | 1.57 | 28.43 | 8.63 | 0.28 | 5.29E+08 | 6.83 | 0.09 |
| 300.200 | -65.85005 | -71.37653 | South | slope | 0.93 | 33.72 | 2151 | 24.00 | 1.60 | 28.73 | 8.28 | 0.15 | 2.54E+08 | 9.83 | 0.05 |
| 300.100 | -66.50618 | -69.87708 | South | shelf | 1.03 | 33.74 | 2149 | 23.02 | 1.59 | 44.15 | 23.68 | 0.75 | 5.75E+08 | 12.27 | 0.21 |
| 300.040 | -66.89105 | -68.92263 | South | coastal | 0.51 | 33.26 | 2095 | 16.39 | 1.32 | 57.59 | 48.92 | 1.97 | 6.35E+08 | 0.68 | 0.61 |
| 200.040 | -67.51042 | -70.59103 | South | coastal | 0.43 | 33.32 | 2095 | 14.88 | 1.09 | 38.72 | 78.27 | 2.29 | 7.72E+08 | 10.98 | 0.76 |
| 200.100 | -67.11405 | -71.53085 | South | shelf | 0.71 | 33.60 | 2139 | 19.98 | 1.50 | 52.09 | 64.60 | 2.01 | 7.62E+08 | 5.52 | 0.64 |
| 200.180 | -66.57926 | -72.73886 | South | slope | 1.09 | 33.72 | 2152 | 23.98 | 1.58 | 30.52 | 97.96 | 0.50 | 6.33E+08 | 12.05 | 0.15 |
| 100.180 | -67.15631 | -74.4745 | Far South | slope | 1.16 | 33.77 | 2157 | 24.71 | 1.65 | 36.76 | 19.40 | 0.55 | 2.78E+08 | | 0.17 |
| 100.100 | -67.70761 | -73.27071 | Far South | shelf | 0.22 | 33.37 | 2128 | 21.83 | 1.54 | 38.25 | 18.63 | 0.42 | 4.14E+08 | 15.50 | 0.15 |
| 100.040 | -68.09312 | -72.38819 | Far South | coastal | 0.22 | 33.12 | 2030 | 5.71 | 0.43 | 39.16 | 162.87 | 6.49 | 1.14E+09 | 74.56 | 2.44 |
| 000.040 | -68.69465 | -74.19341 | Far South | coastal | -1.68 | 33.36 | 2159 | 24.46 | 1.76 | 58.48 | 70.57 | 1.94 | 1.03E+09 | 15.56 | 0.92 |
| 000.100 | -68.27142 | -75.11729 | Far South | shelf | -0.17 | 33.01 | 2121 | 21.51 | 1.51 | 39.00 | 29.35 | 0.57 | 3.95E+08 | 4.22 | 0.24 |
| 000.180 | -67.71603 | -76.29576 | Far South | slope | 0.79 | 33.49 | 2155 | 24.39 | 1.66 | 40.26 | 19.71 | 0.44 | 3.42E+08 | 3.70 | 0.12 |
| -100.180 | -68.25703 | -78.19839 | Far South | slope | 0.63 | 33.62 | 2163 | 24.99 | 1.69 | 36.06 | 14.08 | 0.32 | 3.77E+08 | 4.46 | 0.1 |
| -100.100 | -68.82457 | -77.05133 | Far South | shelf | -0.02 | 33.41 | 2165 | 26.98 | 1.78 | 45.74 | 10.35 | 0.39 | 5.22E+08 | 3.44 | 0.1 |
| -100.040 | -69.25031 | -76.14377 | Far South | coastal | -1.77 | 33.05 | 2117 | 20.88 | 1.45 | 48.24 | 64.72 | 2.16 | 4.45E+08 | 18.54 | 0.79 |
| 200.-040 | -68.03066 | -69.28707 | South | coastal | 1.61 | 33.29 | 2066 | 8.54 | 0.60 | 38.41 | 94.12 | 3.45 | 6.55E+08 | 67.50 | 1.23 |

**Table S2.** Sequencing, assembly, and mapping statistics for metatranscriptomes of each sample.

| Station | Raw sequence reads (million) | Trimmed sequence reads (million) | Number of contigs | Avgerage length (bp) | Maximum length (bp) | Minimum length (bp) | N50 | Mapping efficiency to combined assembly (%) |
|---|---|---|---|---|---|---|---|---|
| 600.040 | 16.3 | 15.9 | 615,687 | 410 | 9704 | 73 | 414 | 81.6 |
| 600.100 | 14.6 | 14.2 | 774,947 | 409 | 7386 | 78 | 400 | 74.9 |
| 600.200 | 14.8 | 14.3 | 729,388 | 386 | 5609 | 73 | 365 | 71.8 |
| 500.200 | 14.2 | 13.8 | 698,566 | 389 | 7552 | 76 | 364 | 70.4 |
| 500.100 | 14.5 | 14.1 | 737,581 | 410 | 6198 | 74 | 394 | 72.1 |
| 500.060 | 13.7 | 13.4 | 592,842 | 425 | 11939 | 75 | 429 | 80.6 |
| 400.040 | 14.2 | 13.8 | 655,372 | 421 | 12191 | 74 | 412 | 73.8 |
| 400.100 | 13.4 | 13.1 | 709,953 | 383 | 6310 | 78 | 357 | 78.0 |
| 400.200 | 14.1 | 13.7 | 632,835 | 379 | 5551 | 74 | 359 | 69.3 |
| 300.200 | 15.0 | 14.6 | 796,624 | 411 | 8601 | 75 | 399 | 73.2 |
| 300.100 | 16.8 | 16.4 | 675,170 | 389 | 6179 | 75 | 376 | 75.8 |
| 300.040 | 14.9 | 14.5 | 714,397 | 373 | 5293 | 73 | 345 | 77.8 |
| 200.040 | 14.6 | 14.2 | 644,421 | 399 | 7363 | 75 | 381 | 70.3 |
| 200.100 | 12.9 | 12.5 | 626,118 | 412 | 6220 | 76 | 404 | 73.3 |
| 200.180 | 14.2 | 13.8 | 648,686 | 402 | 5875 | 73 | 388 | 74.3 |
| 100.180 | 15.8 | 15.4 | 350,342 | 404 | 6473 | 74 | 386 | 78.2 |
| 100.100 | 13.8 | 13.5 | 486,601 | 376 | 4778 | 73 | 359 | 77.0 |
| 100.040 | 13.3 | 12.9 | 660,588 | 405 | 5460 | 73 | 397 | 80.9 |
| 000.040 | 13.2 | 12.9 | 745,478 | 425 | 6544 | 76 | 424 | 72.3 |
| 000.100 | 14.4 | 14.0 | 695,824 | 406 | 7490 | 73 | 388 | 78.2 |
| 000.180 | 101.8 | 99.6 | 792,433 | 406 | 7697 | 73 | 395 | 75.3 |
| -100.180 | 14.4 | 14.0 | 607,556 | 389 | 7008 | 74 | 375 | 73.8 |
| -100.100 | 17.1 | 16.6 | 711,812 | 386 | 7540 | 76 | 362 | 76.1 |
| -100.040 | 14.3 | 13.9 | 350,342 | 373 | 4778 | 73 | 345 | 76.8 |
| 200.-040 | 16.2 | 15.7 | 796,624 | 425 | 12191 | 78 | 429 | 71.5 |
